# Delayed developmental maturation of frontal cortical circuits impacts decision-making

**DOI:** 10.1101/2024.05.24.595609

**Authors:** Kevin Mastro, Wen-Chun Lee, Wengang Wang, Beth Stevens, Bernardo L. Sabatini

## Abstract

In humans, frontal cortical circuits mature in parallel with the development of higher cognitive functions over the course of 15-20 years. In mice, behavior and brain structure change rapidly until reaching sexual maturity (∼8 weeks of age). In contrast, the degree to which frontal cortices and the behaviors they support continue to develop in mice after this period is unclear. Here, we uncover age-related changes in the acquisition and execution of a probabilistic reward reinforcement task. Both young and old mice complete a similar number of trials per session, but 16-30-week-old mice obtain higher reward rates than younger mice. The older mice adjust their behavior more readily after failing to receive a reward, suggesting that age-dependent changes to circuits relevant to forming action-outcome associations, such as those in frontal associative cortices (FACs). Indeed, we identified cell-type and input-specific refinements of FAC circuits over the first 24 weeks of age, including protracted reductions in excitatory drive and increases in inhibition onto pyramidal cells. Dampening inhibitory activity in FAC alters behavior in a manner that counteracts the age-related differences. Together, these reveal an extended period of synaptic maturation in FAC that directly impacts age-related changes in decision-making.

## Introduction

In humans, frontal cortical circuit maturation is a lengthy process that extends into the 2^nd^ and 3^rd^ decade of life and parallels the development of higher cognitive functions, such as goal-directed decision-making[1–4]. Structural changes over this time include synapse formation (birth to 10-12 years old) followed by synaptic pruning and maturation (12 to 20-25 years) and increases in myelination [5–8]. Functional imaging and physiological recordings reveal protracted changes in frontal cortical circuit connectivity and the oscillations they express that correlated with changes in behavior [9–13]. Over the same period, cognitive maturation is reflected in increased executive function and abstract reasoning and reduced impulsivity, which impact goal-directed decision-making [2, 14–18]. Moreover, perturbations of frontal cortical structure are implicated in cognitive deficits found in psychiatric and degenerative diseases [19–25].

In mice, age-dependent maturation of the circuitry of frontal association cortices (FAC) has been described up to ∼8 weeks of life. Structural changes during this time include increased synapse density onto pyramidal neurons and input-specific refinement of synapses onto GABAergic interneurons [26–28]. Consistent with data from humans, lesions to mouse FAC and developmental manipulation of its inputs or local circuitry perturb cognition, including goal-directed decision-making [29–32].

Although the degree of homology of mouse and human FAC is contentious, in both species these areas are innervated by the mediodorsal thalamus (MD) [33, 34]. MD inputs are developmentally regulated, contribute to higher cognitive functions and, in mice, their manipulation affects decision-making [30, 35, 36]. Therefore, refinement of MD input may contribute to protracted development of mouse FAC and associated behavioral changes. Similarly, protracted age-dependent changes in the local GABAergic circuits, whose maturation in sensory cortical areas is thought to underlie the opening and closing of critical periods [37], may contribute to post-adolescent circuit and behavioral refinement. In sensory critical periods, fast-spiking interneurons (FSIs) constrain the developmental period (both onset and offset) and display age-dependent cellular, molecular and functional changes [38–40]. Within frontal cortical circuits, early postnatal increases in inhibition onto pyramidal neurons have been observed and developmental manipulations of FSI activity can either disrupt cognition or is a target for potential therapeutic intervention in mouse model of human disease [26, 32, 41].

Here, we examine the continued development of mouse FAC and behavior in FAC-dependent tasks beyond the age of sexual maturity. We find differences in the acquisition and execution of a FAC-dependent 2-armed bandit task (2ABT) in which thirsty mice must integrate information across trials to optimally achieve water rewards. In parallel, we find changes in both FAC microcircuitry and long-range excitatory inputs after the onset of sexual maturation. Utilizing whole-cell electrophysiology and optogenetics in acute brain slices, we revealed that thalamic, transhemispheric cortical, and local intracortical circuits continue to change after sexual maturity in FAC. At sexual maturation, the relative proportions of inhibition and excitation shift towards a greater influence of inhibitory tone. Furthermore, adult level performance in the 2ABT is sensitive to the degree of inhibition in FAC, such that chemogenetic manipulation of FAC FSIs alters how mice use past experience to select future actions. Together, these data reveal a protracted period of synaptic and behavioral changes associated with the maturation of frontal cortical circuits.

## Results

To determine if FAC-dependent mouse behaviors change after sexual maturity, thirsty mice of a variety of ages were trained in a 2-armed bandit task (2-ABT) that requires them to learn to nose poke (i.e. place their snout) into a variety of ports for potential water rewards. Mice must learn that one port delivers rewards at high probability and the other at low probability. The identity of the highly rewarded port is held constant during “blocks” but switches in a uncued manner after a predetermined number of rewards or trials (**Figure 1a**). Optimizing water acquisition on this task requires the agent to infer the identity of the highly reward port based on reward history and choose this port until the contingencies change. Therefore, at the start of a new block mice must detect that the identity of the highly reward port has changed and adjust their behavior to maximize water acquisition.

**Figure 1.**
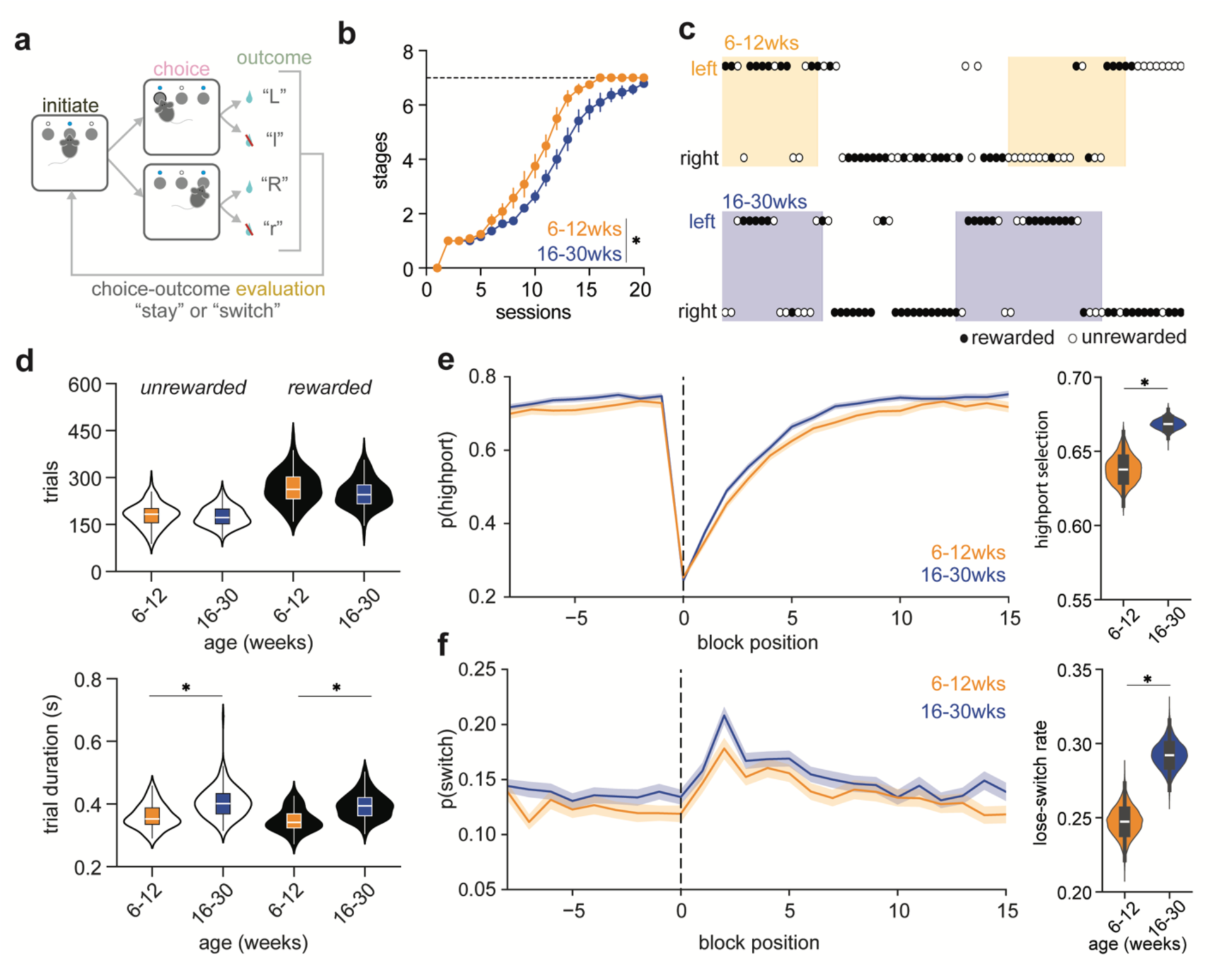
Age related differences in acquisition and execution of a 2-armed bandit task. **a,** Schematic representation of task structure. Mice initiate a trial with a nose poke in the center port followed by a side choice selection whose outcome is probabilistic delivery of a water reward. The identity of the highly rewarded port is held constant for a variable length of rewards during acquisition or trials during evaluation (known as a ‘block’) before the rule is reversed. **b,** Proportion of mice progressing from stage 1 to 7 across daily sessions of training (∼30 mins/session) across two ages (n6-12wks=12 & n16-30wks=18 mice). Two-way repeated measures ANOVA was performed (p=0.001). **c,** Schematic representation of task performance showing that mice select the left (top) or right (bottom) port which is followed by either a rewarded (filled) or unrewarded (unfilled) outcome. Shaded portions show the blocks in which the left port is highly rewarded. **d,** Boxplots superimposed onto violin plots of the number of trials completed per session (top) and trial duration (bottom) that are either rewarded (filled) or unrewarded (unfilled) during the evaluation phase. Student’s T-test comparing rewarded and unrewarded across age groups (p=0.001). **e,** Probability of selecting the highly rewarded port (p(high-port)) at each trial position before (−5 to −1), after (1-12) the rule change (block position=0) (*left*) and violin plot of the bootstrapped probability distributions for selecting the highly rewarded port (*right*) (p<0.0001). **f,** Probability of switching the selected side choice (p(switch)) at each trial position before (−5 to - 1), after (1-12) the rule change (block position=0) (*left*) and violin plot of the bootstrapped probability distributions for switch ports after a loss (*right*) (p<0.0001).

To learn the 2-ABT, mice progress through a curriculum of 6 learning stages that range from easy to the full task. Mice progress from one learning stage to the next after they can acquire a preset number of rewards at that stage difficulty within one session **(Supplementary Table 1)**. Task acquisition is age dependent such that young mice (6-12 weeks old) move through the stages more quickly and learn the full task faster than do older mice (16-30 weeks old) (**Figure 1b**).

After acquiring the full task, mice progressed to the evaluation phase in which the probabilities of receiving rewards are 80% on the highly rewarded port (p(high-port) = 0.8) and 20% on the lowly rewarded port (p(low-port) = 1-p(high-port) = 0.2). Under these uncertain conditions, mice experience both rewarded and unrewarded outcomes at each port throughout each block (**Figure 1c**) and must integrate information across trials to determine the highly rewarded port. During the final five sessions of the evaluation phase, both groups complete the same number of trials and acquire a greater number of rewarded than unrewarded trials but the younger mice complete both rewarded and unrewarded trials significantly faster (**Figure 1d**).

To assess the history dependence of behavior in the task, we examined the probability that the animal selects the highly rewarded port in trials centered around the block boundaries. Towards the end of one block, the behavior of mice in both age groups stabilize such that they select highly rewarded port in ∼2/3 of trials; however, the frequency of selected the highly rewarded port is higher for mice in the older group (**Figure 1e**). As described previously[42], mice exhibit such stochastic port choice, occasionally sampling the lowly rewarded port well into the block, even if the highest reward rate is theoretically achieved by consistently selected the port that is inferred to be highly rewarded. At the uncued block transition, the identity of the highly reward port changes and the probability of selecting it drops for both groups before gradually increasing back to the steady state level (**Figure 1e**). Although mice tend to select the same port on consecutive trials, they do occasionally switch port selections from trial-to-trial. Older mice display greater probability of port switching than younger mice both at steady state and following block transitions (**Figure 1f**). In particular, older mice are more likely to switch ports after failing to receive a reward than are younger mice (i.e., they have a higher lose-switch rate, **Figure 1f**). Together, these data reveal that decision-making processes change during post-adolescent maturation, suggesting that concomitant changes may occur in FAC circuitry.

To determine the developmental profile of FAC during, we examined its circuitry and synaptic properties at different ages, binning data into 4 groups depending on mouse age (2-5, 6-9, 10-15, and 16-30 weeks). We performed whole-cell voltage-clamp recordings from Layer 2/3 (l2/3) pyramidal neurons in the prelimbic (PL) region of the FAC in acute brain slices. An extracellular stimulating electrode was placed 500 µm from pia and the stimulation intensity was adjusted to establish a threshold stimulus that generated a 25-250 pA excitatory response in the voltage-clamped neuron (**Figure 2a-b**). For each cell, synaptic currents were recorded at holding potentials of −70 and 0 mV to isolate excitatory (EPSCs) and inhibitory (IPSCs) postsynaptic currents, respectively, elicited by a range of stimulus strengths relative to the minimal threshold stimulus strength (**Supplementary Figure 2a-d**). Series resistance and membrane capacitance were similar across groups, but membrane resistance is significantly higher at 10-15 and 16-30 weeks relative to other ages (**Supplementary Figure 1e**), suggesting changes in cell size and passive properties across ages.

**Figure 2.**
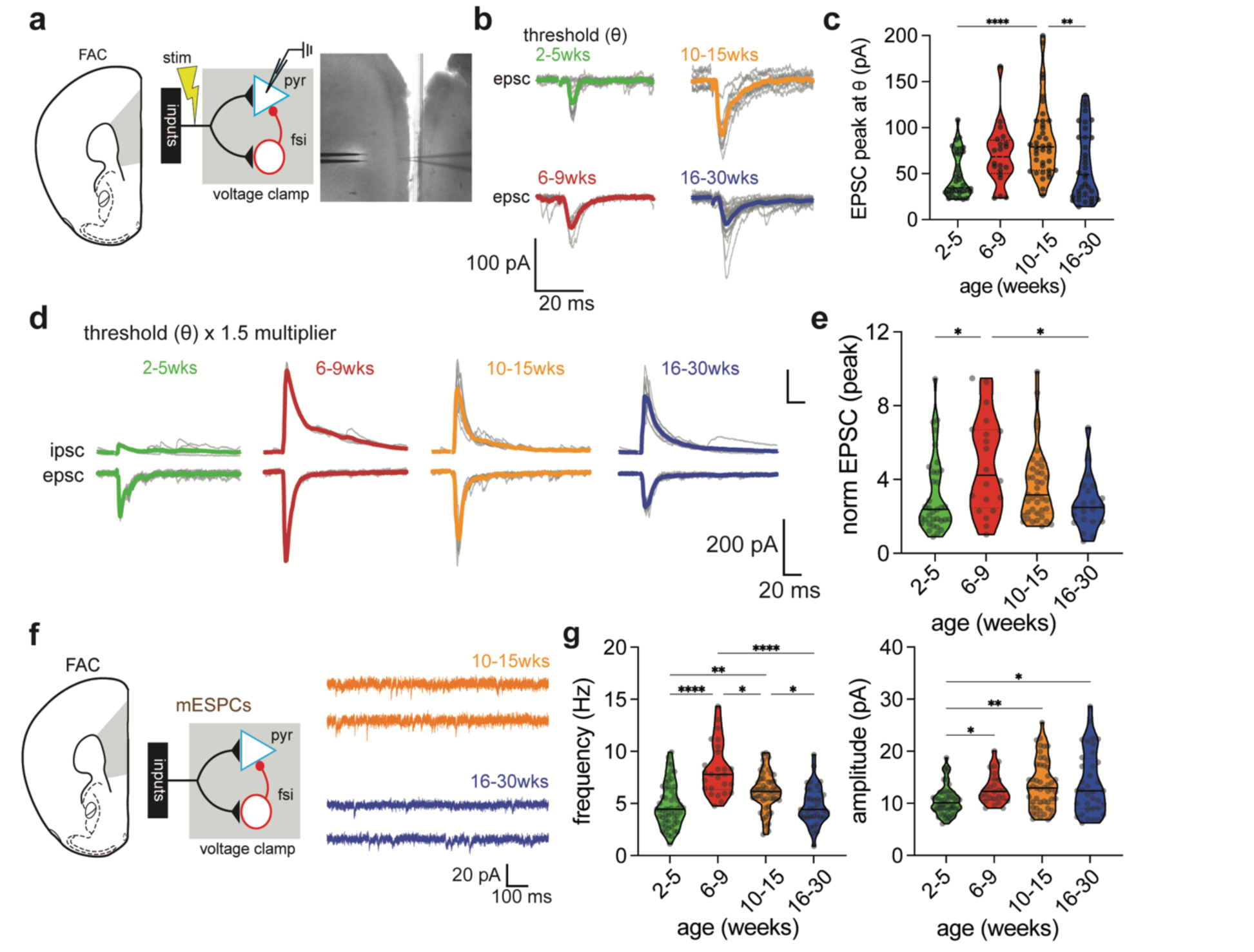
Refinement of excitatory drive onto l2/3 pyramidal (pyr) neurons in the FAC. **a,** Schematic of stimulation and recording configuration for analyzing evoked synaptic currents in l2/3 pyramidal (pyr) neurons with (inset, *left*) low-magnification image of stimulator (two-prong) and recording electrode in frontal association cortex (FAC). The bipolar concentric electrode was placed 500 µm from pia (inset, right) **b,** Representative (gray) and average (overlaid color) of evoked EPSCs at threshold (theta) for an individual l2/3 pyr neuron for each age range. **c,** Evoked EPSC peak amplitude in l2/3 pyr neurons at threshold shown as violin plots (solid line=median, dashed line=upper and lower quartiles) with all points superimposed. For EPSC: (n2-5wks=47, n6-9wks=20, n10-15wks=31, n16-30wks=21 cells). One-way ANOVA was performed and found no difference at threshold (Kruskal Wallis test statistic=5.803; p=0.1216 **d,** Representative (gray) and average (overlaid color) trials of electrically-evoked EPSCs (*Vhold*=-70 mV) and IPSCs (*Vhold*=0 mV) for an individual l2/3 pyr neuron in each age group **e,** EPSC peak amplitudes evoked at 1.5*threshold in each age group shown as violin plots (solid line=median, dashed line=upper and lower quartiles) with all points superimposed (n2-5wks=47, n6-9wks= 20 cells, n10-15wks=31, n16-30wks=21 cells). One-way ANOVA was performed (Kruskal-Wallis statistic=10.49: p=0.0148) with tests for multiple comparisons (* p<0.05) **f**, Schematic of configuration for recording miniature excitatory postsynaptic current (mEPSC) from l2/3 pyr neurons and representative traces of mEPSC recordings from pyr neurons at two of the four ages analyzed. **g**, Frequency (left) and amplitude (right) of mEPSCs in l2/3 pyr neurons from each age group shown as violin plots (solid line=median, dashed line=upper and lower quartiles) with all points superimposed (n2-5wks=54, n6-9wks =33, n10-15wks=51, n16-30wks=36 cells). For frequency: one-way ANOVA was performed (Kruskal Wallis test statistic=40.48; p<0.0001) with tests for multiple comparisons (* p<0.02, ** p=0.0089, **** p<0.0001). For amplitude: one-way ANOVA was performed (Kruskal Wallis test statistic=15.79; p=0.0013) with tests for multiple comparisons (* p<0.03, ** p<0.004)

To compare synaptic properties across the four age groups, we normalized the amplitude of evoked responses to that of the average threshold response per group which, despite our best efforts, varied across the groups (**Supplementary Figure 1a-b**). Comparing the amplitude of current at 1.5 the threshold stimulus strength (1.5x) revealed that this recruited the greatest amount of excitatory drive at 6-9 weeks followed by 10-15 weeks as compared to all other groups (**Figure 2d-e**). The amplitude of inhibitory synaptic currents evoked at this same stimulus strength increased at 6-9 weeks and 10-15 weeks as compared to 2-5 weeks before decreasing again at 16-30 weeks. These age-dependent changes in IPSC amplitudes were consistent across the full range of stimulus strengths (**Supplementary Figure 1c-d**). Together, these data reveal age-dependent reorganization of the frontal cortical circuits but what features of the circuit underlie these changes remain unclear.

To determine whether changes in excitatory connectivity are due to pre-synaptic and/or postsynaptic modifications, we measured the frequency and amplitude of spontaneous miniature excitatory postsynaptic currents (mEPSCs) measured in the presence of TTX (**Figure 2f**). Consistent with anatomical and electrophysiological evidence for increasing spine and synapse density of l2/3 pyramidal neurons over the first 5 weeks of life [26], the frequency of mEPSCs increases from postnatal 2-5 to 6-9 weeks (**Figure 2g**). However, following this peak, mEPSC frequency decreases at 10-15 and 16-30 weeks (**Figure 2g**). In contrast, mEPSC amplitude increases until, and is maintained after, 6-9 weeks (**Figure 2g**). The amplitude of mEPSCs typically reflects post-synaptic properties such as the number of AMPA-type glutamate receptor (AMPAR) per synapse, whereas changes in mEPSC frequency suggest alterations in the number functional AMPAR-containing synapses due to either increases in synapse number, probability of neurotransmitter release, or the fraction of synapses that contain AMPARs (i.e., that are not silent) [43].

Mediodorsal thalamus (MD) and contralateral projecting-FAC (referred to as c-FAC) send dense axonal projections that synapse onto l2/3 neurons within FAC [44]. Disturbances to MD activity through lesions or prolonged inhibition perturb both physiology and behavior [30, 36]. MD innervates FAC early in development and the number of FAC-projecting thalamic neurons decreases over early development in both humans and rodents [45, 46]. Therefore, we hypothesized that changes to MD inputs may contribute to the decreases in the frequency of mEPSCs and amplitude of electrically-evoked responses. To study MD inputs in isolation, we stereotaxically delivered an adeno-associated virus (AAV) encoding channelrhodopsin (ChR2) into MD and, 7-10 days later, prepared acute slices of FAC to examine optogenetically-evoked EPSCs and disynaptic (i.e., feedforward) IPSCs in l2/3 pyramidal neurons (**Figure 3a, Supplementary Figure 2a**). We focused on analysis of mice before (6-9 weeks) the observed reduction in mEPSC frequency or after the reduction had stabilized (16-30 weeks). With the highest laser power, MD-evoked EPSCs are significantly larger at 6-9 than at 16-30 weeks, whereas IPSCs are unchanged (**Figure 3b-c**). These data suggest an overall decrease in the MD projection strength into the l2/3 pyramidal neurons with aging.

**Figure 3.**
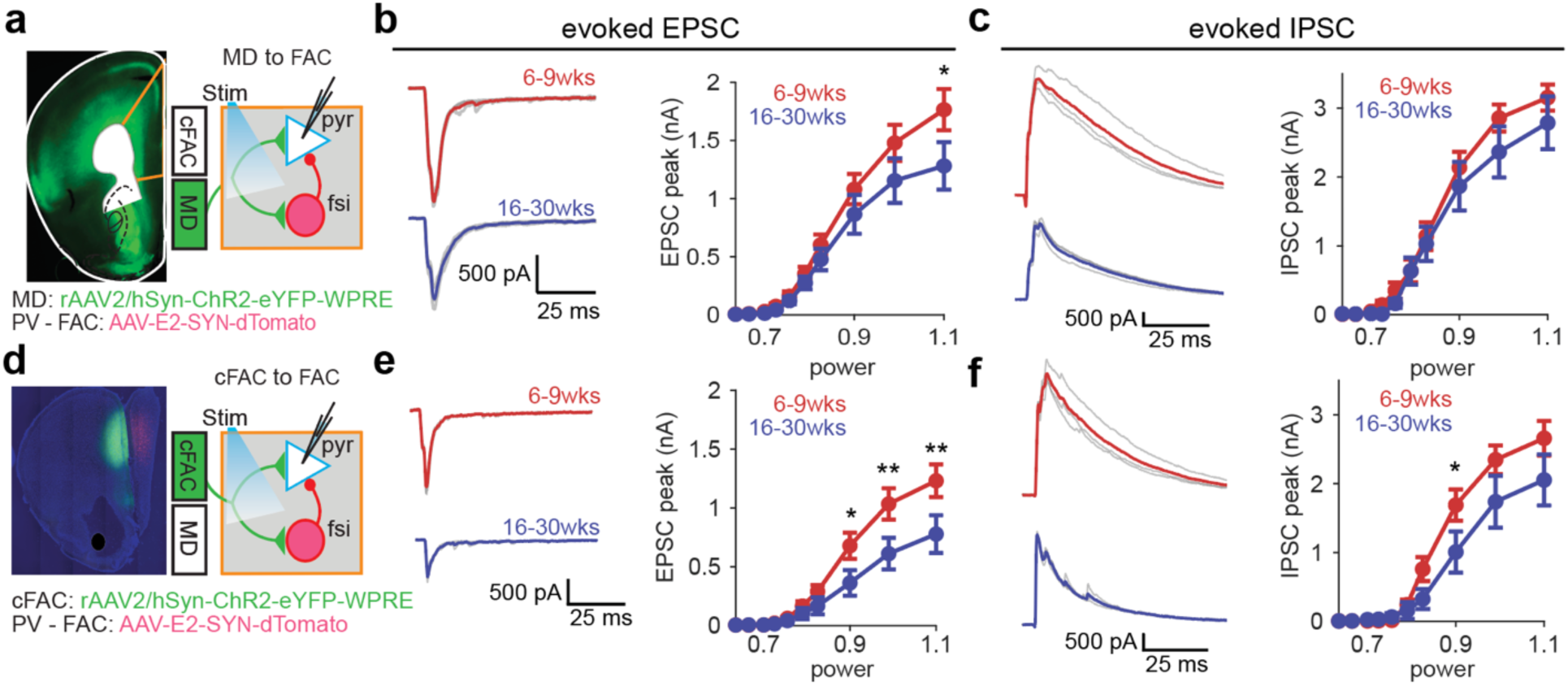
Differential age-dependent refinement of cortical vs. thalamic excitatory drive onto l2/3 pyramidal neurons. **a,** Schematic of experimental preparation showing low-magnification image of expression of ChR2-YFP in mediodorsal thalamic (MD) axons in the FAC (*left*) and recording configuration (*right*). **b**, Representative traces (*left,* average overlaid) (*Vhold*=-70 mV) and peak amplitude of optically-evoked thalamic EPSCs in l2/3 pyrs as a function of laser power in two age groups (n6-9wks=31, n16-30wks=19 cells). Average +/-standard error of the mean shown. Two-way repeated measures ANOVA was performed (p=0.186) with tests for multiple comparisons (* p<0.02) **c**, As in panel **b** but for polysynaptic inhibitory postsynaptic currents (IPSCs) (*Vhold=*0 mV) (n6-9wks=31, n16-30wks=19 cells). Two-way repeated measures ANOVA was performed (p=0.3487). **d,** Schematic of experimental preparation showing low-magnification image of expression of ChR2-YFP in cFAC and expression of the red fluorophore in the FSIs (*left*) and recording configuration (*right*). **e,** As in panel **b** but for optically-evoked cortical inputs to l2/3 pyrs (n6-9wks=19, n16-30wks=17 cells). Two-way repeated measures ANOVA was performed (p=0.053) with tests for multiple comparisons (* p=0.038, ** p<0.002) **f**, As in panel **c** but for polysynaptic IPSCs recorded in l2/3 pyrs and evoked by optical stimulation of cortical inputs (n6-9wks=19, n16-30wks=17 cells). Average +/-standard error of the mean shown. Two-way repeated measures ANOVA was performed (p=0.122) with tests for multiple comparisons (* p=0.0487)

c-FAC sends dense projections to Layer 3 intratelenchepalic (IT) neurons, but the changes in these projections across development have not been explored. Following injections of AAV driving expression of ChR2 in c-FAC, we recorded optogenetically-evoked synaptic currents in l2/3 pyramidal neurons in the contralateral hemisphere (**Figure 3d, Supplementary Figure 2b**). Optogenetic stimulation of inputs from c-FAC revealed a decrease in amplitude of EPSCs across laser powers whereas inhibition was reduced at only one intensity from 6-9 to 16-30 weeks (**Figure 3e-f**).

To investigate whether there synaptic changes are accompanied by age-related changes in intrinsic properties, we obtained whole-cell current clamp recordings from l2/3 pyramidal neurons and examined responses to hyper- and de-polarizing current steps (**Figure 4a-b**). In response to hyperpolarizing currents, neurons from 10-15 weeks old mice had larger voltage changes compared to both 6-9 weeks and 16-30 weeks (**Figure 4b, left)**. We next examined the action potential characteristics at rheobase. The half-width was reduced in both 10-15 weeks and 16-30 weeks compared to 6-9 weeks (**Figure 4c**). Similarly, the afterhyperpolarization (AHP) voltage was more hyperpolarized in 10-15 weeks as compared to the 6-9 weeks and 16-30 weeks (**Figure 4b**). Consistent with this, the maximal firing was slightly increased in this age range relative to both 6-9 and 16-30 weeks timepoints (**Figure 4c**). Overall, intrinsic properties of l2/3 pyramidal neurons are relatively similar across ages.

**Figure 4.**
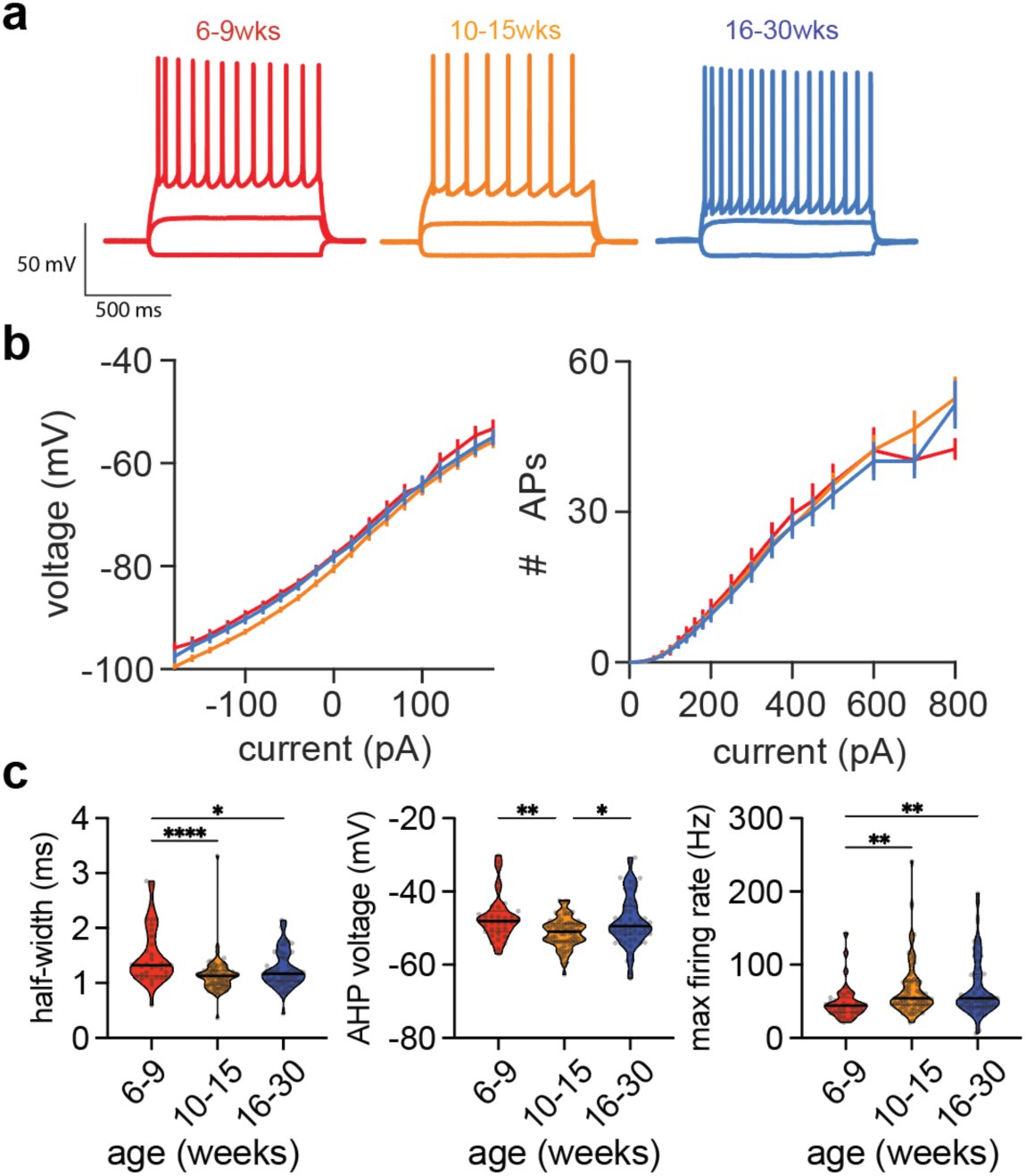
Intrinsic properties and excitability across age. **a,** Representative traces of whole-cell currant clamp recordings of l2/3 pry neurons in the presence of synaptic blockers at each indicated age range. The color code is maintained throughout the figure. **b,** Plateau voltage (*left*) and action potential firing rate (*right*) vs. amplitude of current step (1 s, - 200 to 800 pA) for each age group as indicated in panel **a** **c,** Action potential half-width (*left*), afterhyperpolarization (AHP) voltage (*middle*), and maximum firing rate (*FRmax*, *right*) as a function of current step amplitude for each age group shown as violin plots (solid line=median, dashed line=upper and lower quartiles) with all points superimposed (n6-9wks=35, n10-15wks=71, n16-30wks =55 cells). For half-width: one-way ANOVA was performed (Kruskal Wallis test statistic=19.17; p<0.0001) with tests for multiple comparisons (* p=0.0239, **** p<0.0001). For AHP voltage: one-way ANOVA was performed (Kruskal Wallis test statistic=12.72; p=0.0017) with tests for multiple comparisons (* p=0.0340, ** p=0.0031). For *FRmax*: one-way ANOVA was performed (Kruskal Wallis test statistic=11.18; p=0.0037) with tests for multiple comparisons (* p<0.01)

Fast-spiking interneurons (FSIs), which often express parvalbumin (PV) and cholecystokinin (CCK), typically mediate the fast feedforward inhibition (FFI) in cortex, including that driven by MD inputs [40, 41]. We used either a transgenic mouse (PV-cre) or an adeno-associated virus engineered to express transgenes in PV neurons (E2 enhancer) to express genetically-encoded fluorophores in FSIs for targeted whole-cell recordings. We used the electrical stimulation protocol described above for analysis of EPSCs and IPSCs in l2/3 pyramidal neurons to uncover age-dependent changes in synapses formed onto FSIs (**Figure 5a**). We found significant decreases in the excitatory and inhibitory drive onto superficial, putative FSIs (**Figure 5b, Supplementary Figure 3a-d**) at 16-30 relative to 6-9 weeks. To delineate pathway-specific changes onto FSIs, we again utilized optogenetic-assisted mapping of both MD and c-FAC inputs. MD-evoked excitatory and inhibitory drive was significantly reduced at 16-30 compared to 6-9 weeks (**Figure 5c-d**). In contrast, the c-FAC evoked excitatory and inhibitory drive were unchanged (**Figure 5e-f**). Comparing these results with the concurrently collected data in l2/3 pyramidal neurons shows cell-type and pathway-specific changes over aging (**Supplementary Figure 4**). MD inputs switch from more strongly innervating FSIs to equally targeting FSIs and l2/3 pyramids whereas c-FAC preferentially targets FSIs at both timepoints.

**Figure 5.**
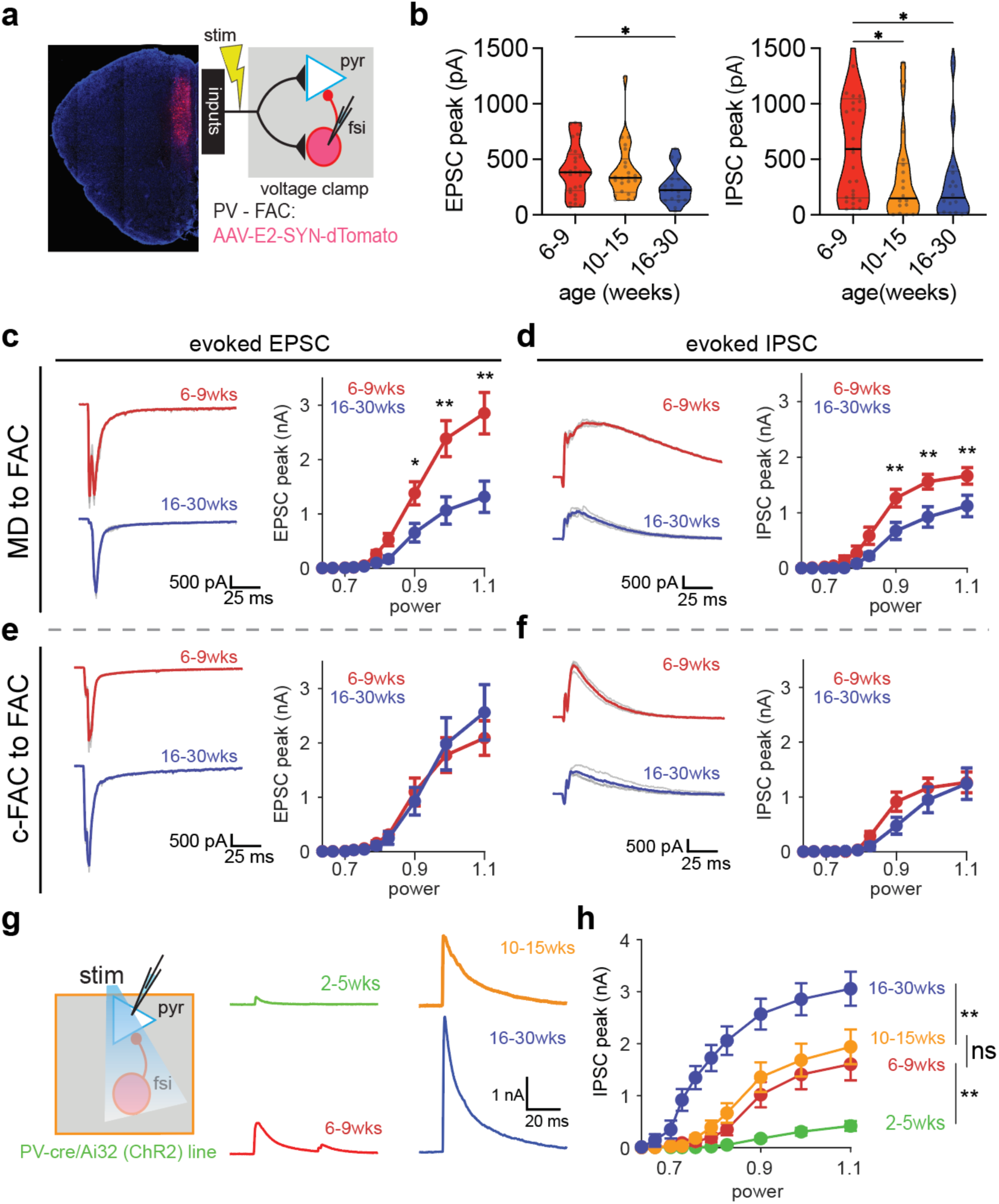
Differential age-dependent refinement of cortical vs. thalamic excitatory drive onto l2/3 fast-spiking interneurons. **a,** Schematic of stimulation and recording configuration for analyzing evoked synaptic currents in fast-spiking interneurons (FSIs) with (inset, *left*) low-magnification image of AAV expression driving fluorophore in putative FSIs in the FAC. The bipolar concentric electrode was placed 500 µm from pia. **b,** Evoked synaptic current peak at 1.5*threshold shown for EPSCs (*left*) and IPSCs (*right*) across ages (n6-9wks=29, n10-15wks=22, n16-30wks=18 cells) shown as violin plots (solid line=median, dashed line=upper and lower quartiles) with all points superimposed. Two-way repeated measures ANOVA was performed (p=0.0328) with tests for multiple comparisons (* p=0.034). **c,** Representative traces (*left,* average overlaid) (*Vhold*=-70 mV) and peak amplitude of optically-evoked thalamic EPSCs in l2/3 pyrs as a function of laser power in two age groups (n6-9wks=15, n16-30wks=20 cells). Average +/-standard error of the mean shown. Two-way repeated measures ANOVA was performed (p=0.002) with tests for multiple comparisons (* p=0.010, ** p<0.0001). **d**, As in panel **c** but for polysynaptic IPSCs recorded in FSIs (*Vhold=*0 mV) (n6-9wks=15, n16-30wks=20 cells). Two-way repeated measures ANOVA was performed (p=0.002) with tests for multiple comparisons (** p<0.0008). **e,** As in panel **c** but for optically-evoked cortical inputs to FSIs (n6-9wks=19, n16-30wks=17 cells). Two-way repeated measures ANOVA was performed (p=0.814). **f**, As in panel **e** but for polysynaptic IPSCs recorded in FSIs and evoked by optical stimulation of cortical inputs (n6-9wks=19, n16-30wks=17 cells). Two-way repeated measures ANOVA was performed (p=0.318). **g,** Schematic of optogenetic stimulation and recording configuration in tissue expressing ChR2 in PV+ FSIs in FAC (*left*) and representative traces (average overlaid) of evoked IPSCs (each cell was held at 0 mV) onto l2/3 pyramidal (pyr) neurons (*right*) in l2/3 pyr neurons at a variety of ages. **h,** Evoked synaptic current peak from low to high laser power in l2/3 pyr neurons across ages (n2-5wks=41, n6-9wks=21, n10-15wks= 18, n16-30wks=9 cells). Average +/-standard error of the mean shown. Two-way repeated measures ANOVA was performed (p<0.0001) with tests for multiple comparisons (NS p=0.372, ** p<0.0001)

To compare changes in excitation and inhibition across cell types, age and experimental paradigm, we calculated an excitation index by dividing the amplitude of the EPSC by the sum of the amplitudes of the EPSC and IPSC (referred to as *R*). For optogenetically-triggered synaptic currents, the profile of *R* over age was input- and cell-type dependent: it was stable for MD-evoked synaptic currents across ages whereas it increased with age for c-FAC-evoked activity in FSIs and not l2/3 pyramidal cells **(Supplementary Figure 4b, d**). For electrically-evoked synaptic currents onto l2/3 pyramidal neurons, IPSCs and EPSCs have similar amplitude at 2-5 weeks, resulting in *R*∼0.5. However, the excitation index decreases at 6-9 weeks before increasing again in the two subsequent time periods **(Supplementary Figure 4e)**. For FSIs, the excitation index was ∼0.5 at 6-9 weeks before first increasing at 10-15 weeks and subsequently decreasing at 16-30 weeks **(Supplementary Figure 4e**). Together, these data reveal cell-type and input-specific changes in synaptic strength and in the relative contributions of excitation and inhibition across age.

To investigate the contribution of FSI-mediated FFI to these synaptic changes, we crossed the PV-*cre* mouse line with the a Cre-conditional ChR2 line (Ai32) to drive expression of ChR2 in all PV cells and measured the evoked inhibitory current onto l2/3 pyramidal neurons (**Figure 5g**). The light-evoked IPSC amplitude increased from 2-5 weeks to 6-9 and 10-15 weeks before increasing greatly by 16-30 weeks (**Figure 5h**). Thus, utilizing a transgenic approach for ChR2 expression, reveals that local inhibitory drive from PV+ interneurons (putative FSIs) undergoes a delayed period of maturation.

Our results demonstrate that FAC circuitry undergoes delayed development maturation, including changes in long-range excitation and local inhibition that, with increasing age, result in greater FSI-mediated inhibition onto l2/3 pyramidal neurons. As these circuit alterations occur concurrently with changes in behavior, we examined if manipulation of inhibition in FAC activity alters behavior in the 2ABT. We used an AAV driving the expression of an inhibitory DREADDs receptor (hM4Di) under the enhancer element (E2) to chemogenetically target putative FSIs (**Figure 6a**). Control mice were handled in the same way with the exception that an E2-AAV expressing a reporter fluorophore was injected. We trained both control and hM4Di expressing adult mice (16-30 weeks) without injection of DREADDs ligand until they completed 15 sessions and achieved expert performance (**Figure 6a**). We subsequently administered the DREADDs receptor antagonist, deschloroclozapine (DCZ, 3 µg/kg) 30 minutes prior to each behavioral session to both sets of mice. DCZ administration perturbed the strategy in the 2-ABT in mice expressing the hM4Di construct, such that they increased their lose-switch rate. This effect was seen by comparing the first two sessions with DCZ administration to the last two without the drug and was specific to hM4Di expressing mice (**Figure 6b**). Thus, acute manipulation of FSIs in FAC alters performance in the 2ABT, consistent with the involvement of activity of these cells in this task. In order to examine the effects of prolonged changes in FSI mediated inhibition on task performance, as occurs with developmental regulation of feed-forward inhibition in FAC, we continue DCZ administration for an additional 8 days. In this chronically manipulated state, hMD4i-expressing mice selected the highly rewarded port less frequently than control mice (**Figure 6c**), recapitulating the behavioral phenotype observed in young mice (**Figure 1d**).

**Figure 6.**
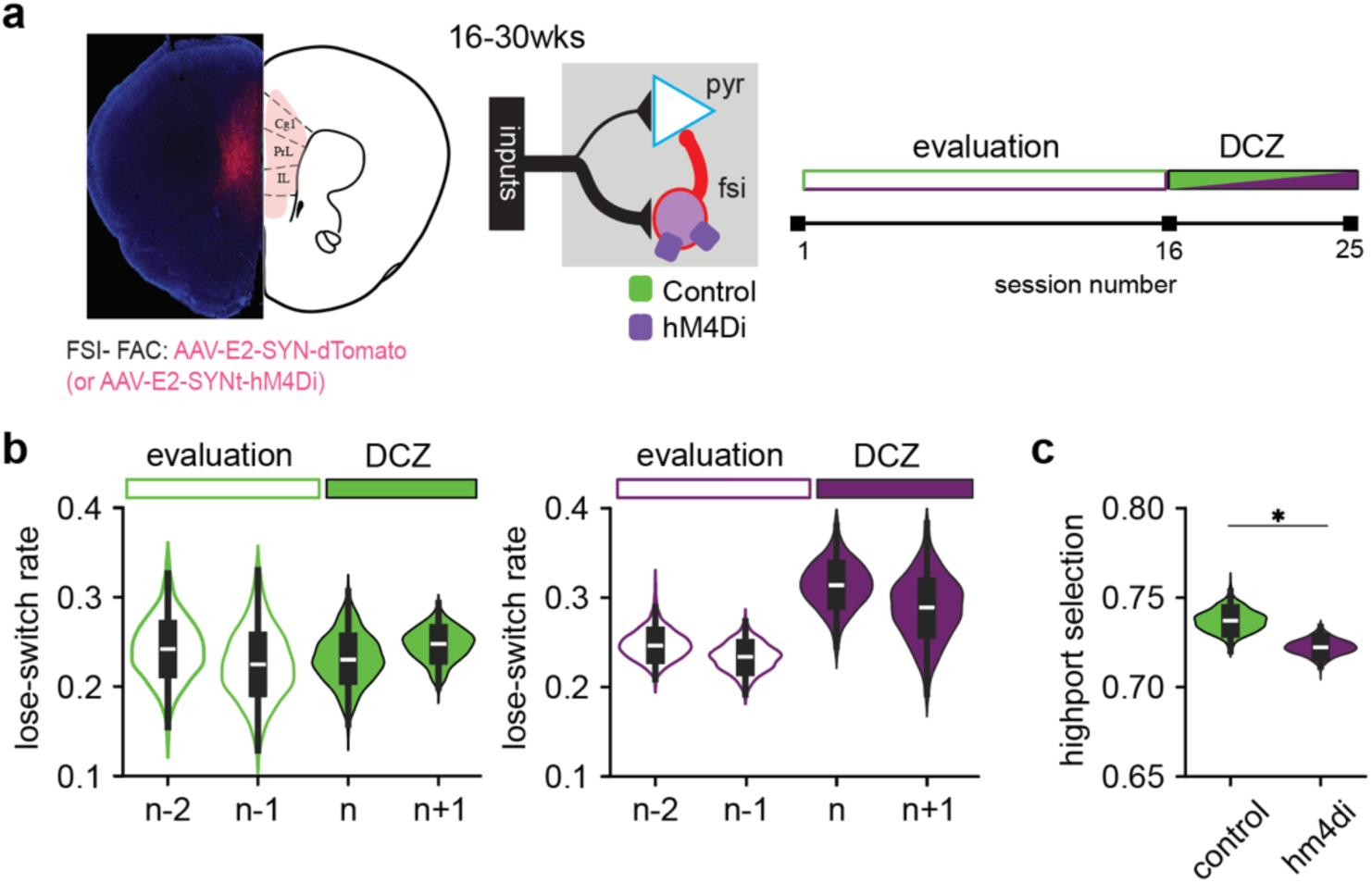
Decreased inhibition reverses age-related changes in 2-armed bandit task. **a,** Representative image (inset, *left*) of frontal association cortical (FAC) expression of AAV driving either fluorophore (dTomato) or inhibitory DREADDs receptor (HM4Di) under the synapsin (SYN) promotor and with the putative fast spiking interneuron (FSI) enhancer (E2). Schematic of receptor expression (inset, *right*) in FSIs and not in pyramidal neurons. Schematic of experimental timeline (*right*): 15 sessions of evaluation without drug administration followed by 10 sessions with administration of DCZ (1-3 mg/ml) via intraperitoneal injection 30-minutes prior to each session. **b,** Bootstrapped distributions of the lose-switch rate for control fluorophore (*left*) and hM4Di (*right*) expression animals before (unfilled; n-2, n-1) and after (filled; n, n+1) the start of the sessions with DCZ administration (session n). **c,** Bootstrapped distribution of highport selection rate in the final five sessions of DCZ administration in hM4Di and control flourophore expressing mice.

## Discussion

We find that the mouse frontal cortical circuits undergo a protracted period of maturation characterized by changes in excitatory and inhibitory inputs onto l2/3 pyramidal neurons and FSIs. These changes reflect pathway-specific refinements of thalamic and intracortical innervation of these two cells types, as well as changes in local inhibitory circuits, during the first 6 months of life. Behaviorally, we find age-related differences in both the acquisition and execution of a probabilistic 2-ABT over the same period such that 6-12 week old mice acquire the task more quickly than their 16-30 week old counterparts. After task acquisition, the older mice are more likely to both select the highly reward port and to switch between ports trial-to-trial, reflected differences in task execution. Chemogenetic inhibition of FSIs in 16-30 week old mice acutely and chronically alters behavior in the 2ABT. Furthermore, chronic inhibition results in task performance in older mice that resembles that of 6-12 weeks old mice. Thus, refinement of FAC inhibition contributes to age-related changes in behavior and to the prolonged development of the FAC circuits.

### Cortical inhibition regulates plasticity and cognitive behaviors

Periods of heightened plasticity, known as critical periods, have been extensively studied in sensory areas where modulation of inhibition is postulated to open and close plasticity windows [39]. Furthermore, experience-dependent refinement of GABAergic circuits is coordinated across multiple inhibitory cell-types and is necessary to establish the precise wiring of cortical circuits. Specifically, FSIs and low-threshold spiking interneurons (LTSI) establish their connectivity at different rates and their post-natal maturation has been observed across multiple sensory areas [47–50]. We show with electrical stimulation that FAC inhibitory circuits continue to refine in a complex manner over a prolonged 30-week period, with increases in evoked inhibition at 6-15 weeks of age relative to younger and older ages. In contrast, selective activation of PV-mediated inhibition shows that the inhibition of l2/3 pyramidal neurons by FSIs increases steadily across this same period. These data suggest that continued maturation of other inhibitory circuits also occurs, including potentially that mediated by LTSIs.

Similar changes occur during the protracted development of human frontal cortex, which includes decreases in excitation and increases in GABA-mediated inhibition until excitatory and inhibitory ratios stabilize in adulthood [51, 52]. Greater inhibitory tone has been postulated to contribute to the refinement of higher cognitive functions. Nevertheless, the specific cellular and synaptic changes that results in the correct excitatory and inhibitory drives at the levels of individual cells and the whole circuits are not fully understood.

### Age impacts decision making

Following sensory critical periods, the brain continues to develop and refine into adulthood, which impacts decision making in humans and rodents [2, 54]. In these species, the maturation of frontal cortical circuits has been linked to behavioral changes and lesions of specific frontal circuits perturb decision-making. From an evolutionary perspective, the frontal cortex has undergone the greatest expansion within mammals and is most complex and enlarged in primates. Although rodent frontal cortex lacks the regionalization and some of the anatomical attributes (e.g., granularity) found in humans, there are anatomical and functional parallels. First, rodent frontal cortex has a similar pattern of connectivity with thalamus and other subcortical regions as primate frontal cortex. Second, lesions to distinct regions perturb decision-making. Lastly, consistent with observations in humans, changes to frontal cortical structure and function are implicated in the modulation of decision-making during development, diseases and degeneration.

The data presented here illuminate the complex changes that occur in one part of mouse FAC circuitry over a protracted developmental period and that influence the relative activation of inhibitory and excitatory cortical neurons. However, they likely comprise only a small part of the circuity changes that occur over this period. In addition to the inputs from MD and c-FAC, FAC is innervated by other cortical areas, ventromedial thalamus, amygdala, hippocampus and VTA. Moreover, the FAC contains additional laminar organization and a diverse set of interneurons which likely also change over this extended period. In addition, both the refinement and action of neuromodulators, including dopamine, likely contribute to post-sexual maturity circuit development, as occurs in the maturation of human FAC and reward processing systems. Therefore, cell-type and input-specific chemogenetic manipulation of FAC FSIs, such as done here, is unlikely to fully revert the behavior of an aged animal to that of a younger one. Nevertheless, it does demonstrate the sufficiency of changes in the activity or synapses of specific interneuron population, such as we find occurs during protracted FAC development, to influence complex behaviors. Our data reveal that blunting FAC inhibition does not influence all aspects of 2ABT behavior but does modulates switching behavior following a negative outcome. In future experiments, will address the contribution of other circuit elements to age-dependent changes in cognitive behaviors in mice.

### Linking synapses to behavior: implications for aging

The synaptic and circuit changes demonstrated here are likely driven by age-dependent transcriptional, neuromodulatory, and experience-dependent processes in both neuronal and non-neuronal cells. Microglial and astrocytic pruning of synapses are key candidates for the mechanisms of developmental circuit sculpting. In addition, oligodendrocytes contribute via post-adolescence myelination including in frontal cortex (FC) which is one of the last brain regions to myelinate. These programs likely also contribute to neuropsychiatric disease, including schizophrenia and other disorders that typically manifest during the transition from adolescence to adulthood and whose risk factors include genes involved in synaptic, glial, and neuroimmune pathways [55–57]. However, further characterization of developmental and disease associated circuit changes, as well as quantitative models in which to examine the functional implications of these alterations, are necessary to understand their full contribution to normal aging and disease progression.

## Methods

All procedures were performed in accordance with the guidelines established by the National Institutes of Health and with approval from the Harvard Medical School Institutional Animal Care and Use Committee.

### Animals

Male mice ranging from 2-30 weeks old on the C57BL/6J (Strain #:000664) background from Jackson Laboratory were employed for all studies. To target a fluorophore or opsin to parvalbumin-positive interneurons, we crossed the PV-Cre (Strain #: 017320) to Ai14 (dTomato, Strain #:007914) or Ai32 (ChR2: Strain #:024109) mouse lines, respectively.

### Viruses

Recombinant adeno-associated viruses (AAVs with stereotypes 2/9, 5 and PHP.eB) were used to identify, monitor or manipulate cell-types utilized viral strategies in the C57BL6/Jax (Strain #:000664) mouse cohort via neuronal or PV-specific enhancer elements[58]. Most viral plasmids can be found at Addgene: AAV5-CamKII-GCaMP6f-WPRE-SV40 (no. 100834-AAV5), AAV9-hSyn-hChR2(H134R)-EYFP (no. 26973-AAV9), and AAV-PHP.eB-S5E2-dTom-nlsdTom (no. 135630-PHPeB). Some custom plasmids were packaged commercially (WZ Biosciences): AAV-PHP.eB-S5E2-hM4D(Gi)-mCherry. The E2-enhancer element provided directed expression in putatively parvalbumin-positive interneurons in frontal association cortex[58]. Upon delivery, viruses were separated into single-use aliquots (3-5 uL) and stored in the −80°C. Injection volumes ranged from 200-250nL and titers were adjusted to approximately 10^11^ to 10^12^ genomic copies per ml.

### Surgery

Stereotaxic injections were performed similarly to previously reported studies[59]. Briefly, mice were anesthetized deeply using isoflurane before being placed in a stereotaxic frame where body temperature and level of anesthesia were monitored throughout the procedure. For all surgeries, the skull was exposed and the distance from bregma to lambda was used to both level and normalized for changes in skull size over the lifespan (normalization factor = measured distance (mm) /4.2 mm). Coordinates were established from bregma and a drill was used to make small craniotomies for viral injections. We used the following coordinates used for each brain area: prelimbic region of the FAC (Area 24: A/P = +1.85, M/L = +/-0.40, D/V = −1.85) and mediodorsal thalamus (MD: A/P = −1.3, M/L = +/-0.55, D/V = −3.05). Pulled electrodes containing the virus would be slowly lower to the dorsal/ventral coordinate and allowed to sit for 5 minutes. Infusion of the virus took 4-5 minutes at a rate of 50 nl/min and was followed by an additional 5 mins to counteract any flow up the injection site. After completing the injections, skin was returned and sutured and ointment was used to prevent any infection and limit inflammation.

### Behavior

The two-armed bandit task (2-ABT) was consistent in both box and behavioral design as previous work[42, 59]. Briefly, mice were water regulated to 1-2 ml/day until they maintained ∼80% *ad libitum* weight. On training days, mice were placed in a 4.9” x 6” acrylic chamber that contained three custom nose ports with an infrared-beam sensor (full design plans can be found on <u>“Open Behavior”</u>).

To facilitate the acquisition of the task, mice progressed through a series of stages. As shown in **Supplementary Table 1**, parameters of the tasks were shifted until achieving the ideal task structure where decisions are made within 2 seconds (“Decision Window”) where the probability of the highly rewarded port (“p(high port)”) was 0.8, probability of the low rewarded port (“p(low port)”) was 0.2 with a variable blocklength of 10-30 trials. For example, Stage 1 rewards the agent for any pokes in the right and left ports. This stage is necessary to acquire the relationship between a nose poke action and the reward. Stage 2 requires the agent to poke in the center port followed by side port poke and reward. This center to side association is strengthened through a combination of decreasing the amount of time allowed between the two pokes (Stage 2-5) followed by a reduction in the length of a given block (Stage 6). To complete a stage, animals had to successfully acquire 100 rewards in a given session for each of stages 1-5 and 200 rewards for Stage 6. Upon successful acquisition of the task, mice were progressed to stage 7 where they completed 8-12 sessions of the probabilistic task. Behavioral training was completed during the dark phase of the mouse sleep/wake cycle and were run under no ambient light.

### Behavioral Analysis

All metrics were based on completed trials where mice initiated a center nose poke followed by the selection of a right or left nose poke. To observe differences in training, the number of days to achieve criterion on any given stage was plotted.

To evaluate the performance on the probabilistic task, the probability that a mouse selects the highly reward port can be calculated relative to the trial position in the block. At ‘Block Position’ zero, the highly rewarded port switches to the other port (From left to right, or right to left) and the mouse must adapt their behavior to track the change in the task. The action of the mouse can be represented by either the probability of correctly choosing the highly-rewarded port (p(high port)) or the probability of switching (p(switch)) which would be expected to increase after block position 0.

We employed a bootstrapping approach to estimate the means and standard deviations of several behavioral metrics across conditions or age groups. For each metric, data was resampled with replacement 500 times to generate bootstrapped distributions. To quantify the differences between conditions, we calculated the delta (Δ) values by subtracting the bootstrapped estimates of the control group from those of the treatment group for each iteration (p120-180 vs p60-100 or Control vs hM4Di, respectively). This generated a distribution of delta values, representing the pairwise differences between the two groups.

### Slice physiology preparation

Coronal sections (250-300 um thickness) were prepared from brains of 5-28 weeks-old mice. First, mice were transcranially perfused with ice-cold ACSF solution (in mM): 125 NaCl, 2.5 KCl, 25 NaHCO3, 2 CaCl2, 1 MgCl2, 1.25 NaH2PO4 and 11 glucose (300–305 mOsm kg−1). Slices from the anterior portion of the mouse brain that contained prelimbic region of the FAC were prepared on a Leica VT1200 vibratome in the same ice-cold ACSF solution followed by a 5-10 min incubation period at 34 °C containing a choline-based solution (in mM): 110 mM choline chloride, 25 mM NaHCO3, 2.5 mM KCl, 7 mM MgCl2, 0.5 mM CaCl2, 1.25 mM NaH2PO4, 25 mM glucose, 11.6 mM ascorbic acid and 3.1 mM pyruvic acid. The choline-based incubation is followed by an additional 15-minute incubation in ACSF at 34 °C. After the second incubation, slices were moved to room temperature and maintained for the length of the recording. All solutions were carbogenated at all stages of tissue preparation and data collection.

### Slice physiology collection and analysis

#### General recording principals

Recordings were made at 32-34°C in carbogenated ACSF (in mM): 125 NaCl, 26 NaHCO3, 1.25 NaH2PO4, 2.5 KCl, 12.5 glucose, 1 MgCl2, and 2 CaCl2. Data was collected on and a low-pass filter was applied (MultiClamp 700B amplifier; Molecular Devices) and then digitized on a National Instruments acquisition board. Data was acquired via a custom version of ScanImage written in MATLAB. Data was analyzed offline using custom code written in Matlab or Python. Electrodes were made from borosilicate glass (pipette resistance, 2– 4 M). Series resistance was monitored throughout the voltage clamp recordings and prior to the start of all current clamp recordings and cells were excluded if series changed greater than 20% of the initial baseline.

#### Voltage clamp recordings

utilized a cesium-based internal (in mM): 135 CsMeSO3, 10 HEPES, 1 EGTA, 3.3 QX-314 (Cl^−^ salt), 4 Mg^−^ATP, 0.3 Na-GTP, 8 Na2-phosphocreatine, pH 7.3 adjusted with CsOH; 295 mOsm kg^−^1). For the external ACSF solution for the electrical and optogenetic experiments, no blockers were used in the external ACSF. For miniature excitatory postsynaptic (mEPSC) experiments, external ACSF contained 1 μM TTX and 10 uM CPP. Voltage-clamp recordings were filtered at 2 kHz and digitized at 10 kHz.

For mEPSC recordings, cells were held at −70 mV and monitored for 5-10 mins. Recording epochs with stable baselines and access resistance were included for analysis. To measure the frequency and amplitude of the excitatory events, we utilized custom MATLAB scripts adapted [60]and applied previously[61].

For electrical stimulation, a bipolar concentric electrode was positioned 500 um from the top of pia and recordings targeted superficial pyramidal neurons and putative fast-spiking interneurons labeled by either viral or transgenic approaches. As described previously[62], threshold (theta) was established by tuning the electrical stimulation (1 pulse @ 10 ms) until a 20-150 pA (peak magnitude) excitatory postsynaptic current was present at the −70 mV holding voltage. The threshold was then multiplied by 1.1-1.5 and responses were recorded at both −70 mV and 0 mV for each neuron. Each stimulation intensity was collected five times with a 30 second inter-stimulation interval. For optogenetic stimulation, laser power was calibrated to the same level of laser intensity. Stimulation (1 pulse @ 3 ms) were taken at varying laser intensities and were randomly interspersed. Each laser intensity was collected three times with a 30 second inter-stimulation interval. Like electrical stimulation, cells were held at −70 mV (to isolate excitatory drive) followed by 0 mV hold (to isolate inhibitory drive). To understand how each stimulation impacted the local network, the current response for each stimulation was integrated over a 20 ms window of time (Area under the curve). To isolate the impact of monosynaptic drive, peak magnitude was analyzed within a short latency activation (<5 ms). For statistical testing, multiple unpaired t-tests adjusted for multiple comparisons were performed and both the p-value and q-value (adjusted for False Discovery Rate) are reported.

#### Currently clamp recordings

utilized a potassium-based internal: 130 KMeSO3, 3 KCl, 10 HEPES, 1 EGTA, 4 Mg-ATP, 0.3 Na-GTP, 8 sodium phosphocreatine (pH 7.3 adjusted with KOH; 295 mOsm). For recordings of intrinsic excitability, 10 uM gabazine 10 uM NBQX and 10 uM CPP were included in the bath to block fast inhibitory and excitatory transmission, respectively. Data was filtered at 10 kHz and digitized at 40 kHz. Each step included a −100 pA step for 100 ms for the calculation of input resistence. For current-voltage relationship, current steps were applied for 1000 ms from −200 to 200 pA in 20 pA steps. For current-firing rate relationship, additional steps were applied from 200-1500 pA in 50-100 pA steps. Cells were monitored for inactivation and steps following inactivation were excluded from analysis.

### Behavioral Chemogenetics

Mice underwent surgery and were randomly assigned to receive injection of HM4Di or mCherry viral vectors. Following surgical recovery and water regulation, mice completed acquisition (Stages 1-6) and 15 sessions of the probabilistic (Stage 7) of the 2-ABT. After which animals were administered the DREADDs activator deschloroclozapine (DCZ: 1-3 mg/ml) via intraperitoneal injection 30-minutes prior to completing the task for 10 sessions [63]. Experimenters were blind to experimental conditions during data acquisition.

### Histology and Imaging

Mice were deeply anesthetized with isoflurane and perfused transcardially with 20 ml of PBS followed by 10 ml of 4% paraformaldehyde (PFA). Brain was excavated and placed overnight in 4% PFA at 4°C. Brains were transferred to 30% sucrose for equilibration and stored at 4°C until falling to the bottom of the container. Using dry ice, brains were frozen in an optimal cutting temperature compound (O.C.T.: Tissue-Tek) and cut on the cryostat (50 um slice). DAPI mounting medium (Vectashield, H-1200) was used to preserve fluorescence quality and was imaged immediately on a widefield microscope at 10x objective. For viral verification, no enhancement of the expression was necessary.

### Statistics

Statistical analysis was completed using custom MATLAB and Python scripts and Prism (GraphPad; Version 10).

## Supplementary Figures

**Supplementary Figure 1.**
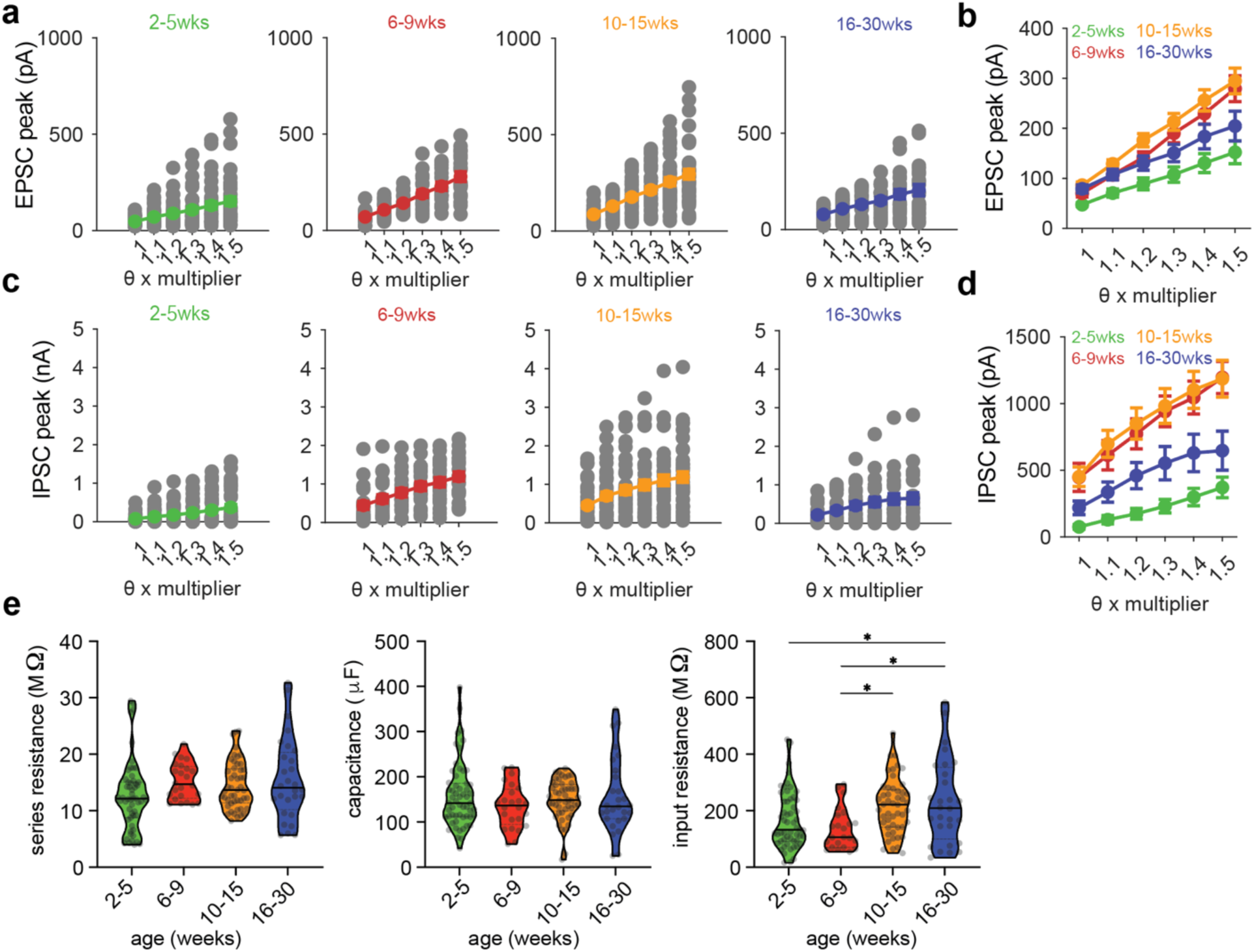
Protracted development of excitatory and inhibitory drive onto l2/3 pyramidal (pyr) neurons in the frontal association cortex (FAC) **a,** Evoked synaptic current peak from threshold to threshold x 1.5 multiplier shown for excitatory (EPSC) postsynaptic currents onto l2/3 pyr neurons (grey=individual cells overlaid with average) across ages (n2-5wks=47, n6-9wks=20, n10-15wks=31, n16-30wks=21 cells). **b,** Averaged evoked synaptic current peak from threshold to threshold x 1.5 multiplier across ages. Average +/-standard error of the mean shown. Two-way repeated measures ANOVA was performed (p=0.052) with tests for multiple comparisons (* p=0.007). **c,** As in panel a but for polysynaptic inhibitory postsynaptic currents (IPSCs) onto l2/3 pyr neurons (*Vhold=*0 mV). **d,** As in panel b but for the average polysynaptic inhibitory postsynaptic currents (IPSCs) (*Vhold=*0 mV) onto l2/3 pyr neurons (n2-5wks=47, n6-9wks=20, n10-15wks=31, n16-30wks=21 cells). **e,** Series resistance (left), capacitance (μF) (middle) and Input resistance (right) shown as violin plots (solid line=median, dashed line=upper and lower quartiles) with all points superimposed (n2-5wks =54, n6-9wk=33, n10-15wks=51, n16-30wks=36 cells). **All cells included from electrical stimulation experiment, even those where stimulation led to greater than 250 pA excitatory response**. For series resistance: one-way ANOVA was performed (F-statistic=2.37; p=0.0790). For capacitance: one-way ANOVA was performed (F-statistic=0.9209; p=0.4428). For input resistance: one-way ANOVA was performed (F-statistic =5.168; p=0.0020) with tests for multiple comparisons (* p<0.02, ** p<0.3 ** p<0.004).

**Supplementary Figure 2.**
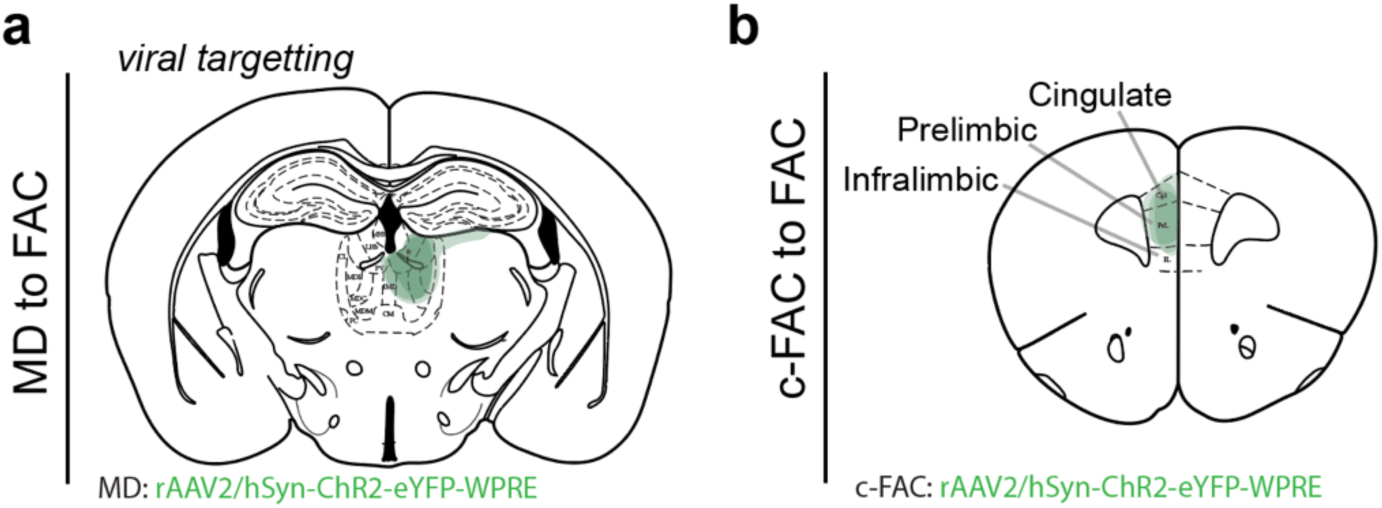
Targeting location for optogenetic-assisted mapping of mediodorsal thalamic and contra-frontal cortical inputs. **a,** Representative viral targeting location across 3 animals in the mediodorsal thalamus (MD), injected with the AAV driving expression of channelrhodopsin (ChR2) in neurons (human synapsin (hSyn) promoter) **b,** Representative viral targeting location across 3 animals in the contralateral-frontal association cortex (c-FAC), injected with the AAV driving expression of ChR2 in neurons (hSyn promoter)

**Supplementary Figure 3.**
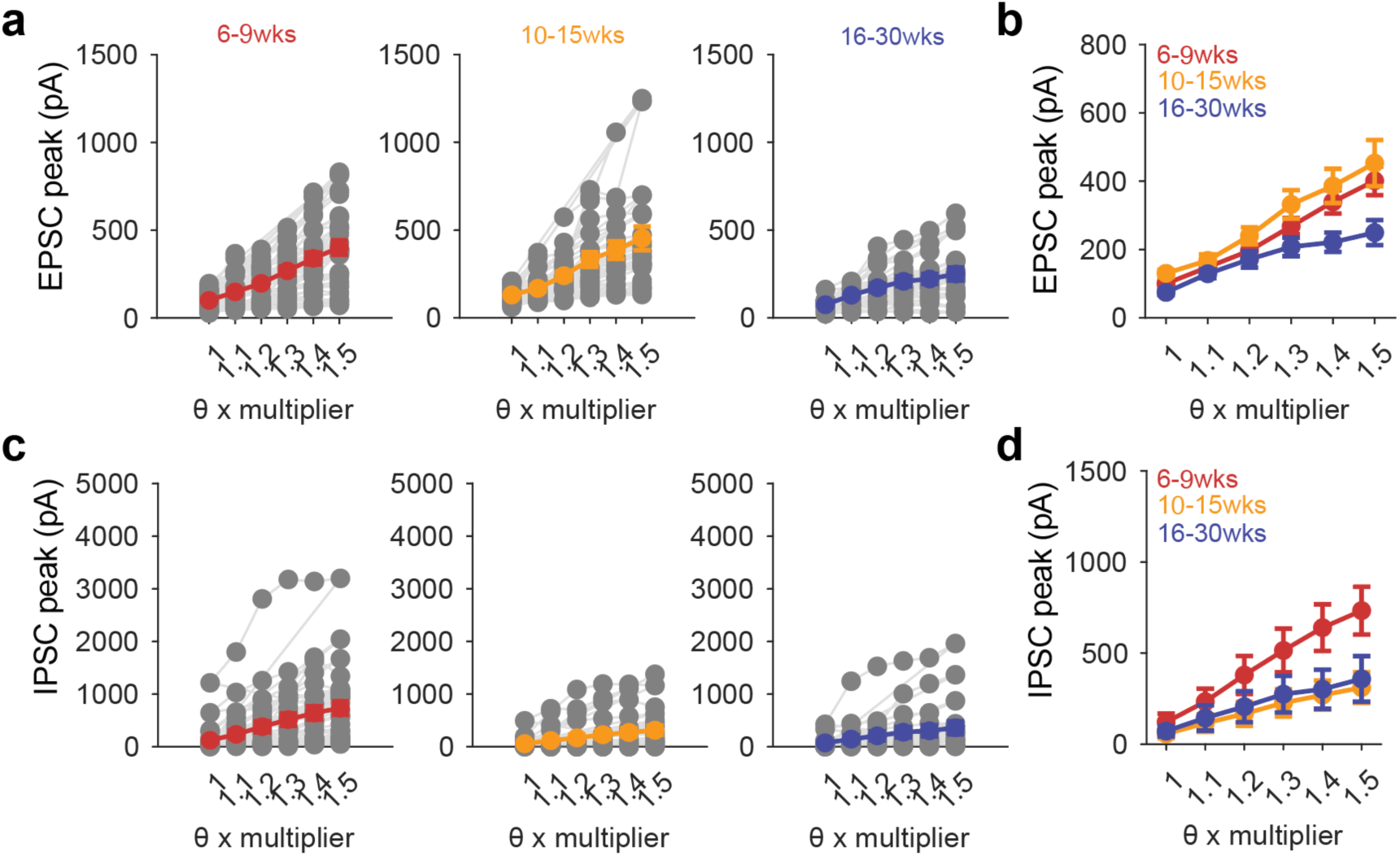
Protracted development of excitatory and inhibitory drive onto fast-spiking interneurons (FSIs) in the frontal association cortex (FAC) **a,** Evoked synaptic current peak from threshold to threshold x 1.5 multiplier shown for excitatory (EPSC) postsynaptic currents onto FSIs (grey=individual cells overlaid with average) across ages (n6-9wks=29, n10-15wks=22, n16-30wks=18 cells). **b,** Averaged evoked synaptic current peak from threshold to threshold x 1.5 multiplier across ages. Average +/-standard error of the mean shown. **c,** As in panel a but for polysynaptic inhibitory postsynaptic currents (IPSCs) onto FSIs (*Vhold=*0 mV). **d,** As in panel b but for the average polysynaptic inhibitory postsynaptic currents (IPSCs) (*Vhold=*0 mV) onto FSIs (n6-9wks=29, n10-15wks=22, n16-30wks=18 cells).

**Supplementary Figure 4.**
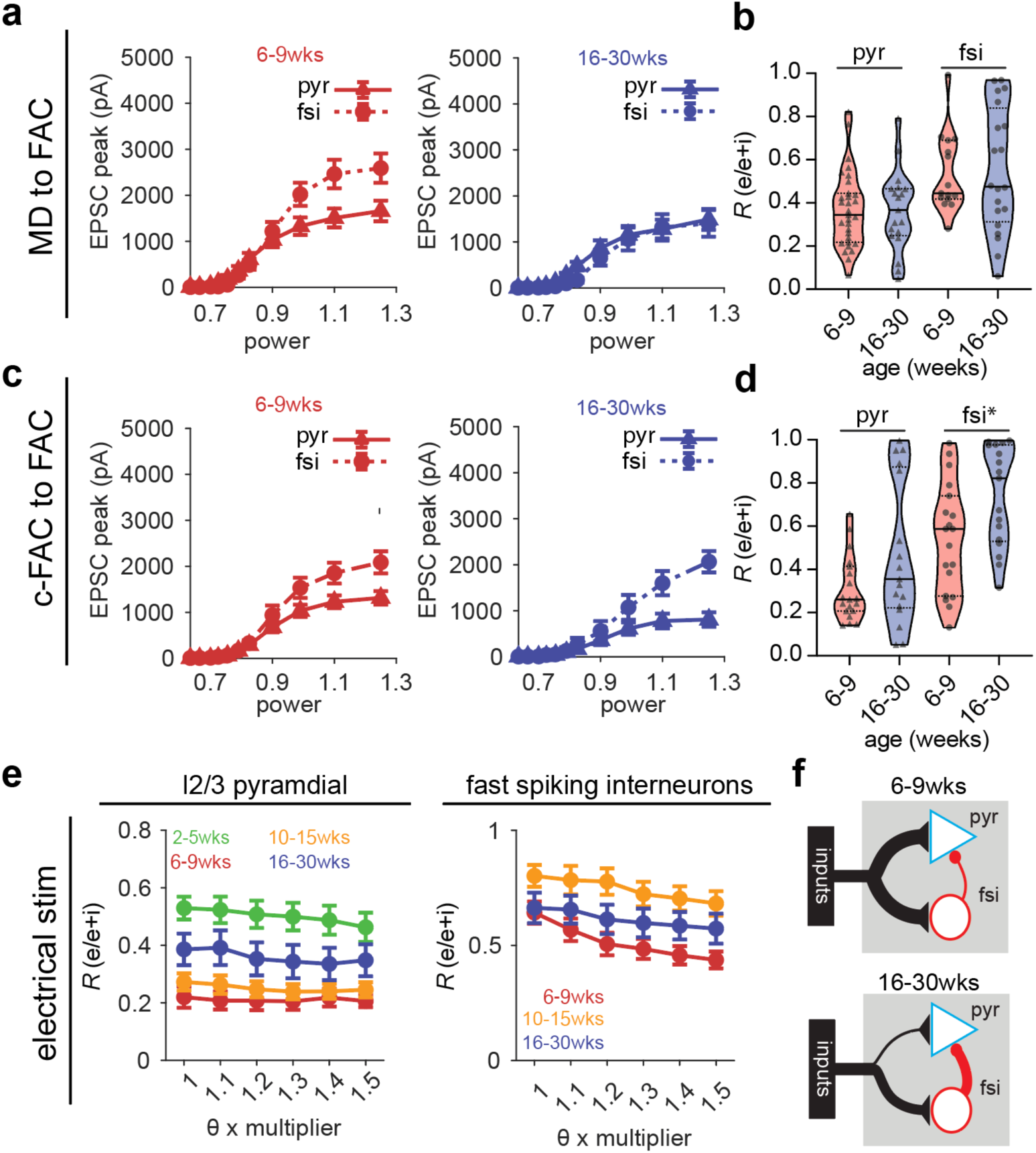
Protracted refinement of excitatory/inhibitory balance drive onto l2/3 pyramidal neurons and fast spiking interneurons across experimental paradigms. **a,** Optically-evoked mediodorsal (MD) thalamic EPSCs onto l2/3 pyramidal (pyr) neurons and fast-spiking interneurons (FSIs) as a function of laser power at 6-9 weeks (left) and 16-30 weeks (right). Average +/-standard error of the mean shown. **b,** Fraction of EPSC over total synaptic inputs (EPSC+IPSC) onto l2/3 pyr (left) and FSIs (right) vs. the optogenetic stimulation of MD inputs across the two ages shown as violin plots (solid line=median, dashed line=upper and lower quartiles) with all points superimposed (pyr: n6-9wks=31, n16-30wks=19 cells & fsi: n6-9wks=15, n16-30wks=20 cells). **c,** As in panel a but for optically evoked contracortical (c-FAC) inputs onto currents l2/3 pyr neurons and FSIs at 6-9 weeks (left) and 16-30 weeks (right). **d,** Fraction of EPSC over total synaptic inputs (EPSC+IPSC) onto l2/3 pyr (left) and FSIs (right) vs. the optogenetic stimulation of c-FAC inputs across the two ages shown as violin plots (solid line=median, dashed line=upper and lower quartiles) with all points superimposed (pyr: n6-9wks=19, n16-30wks=17 cells, fsi=(n6-9wks=19, n16-30wks=17 cells). **e,** As in panel b but for electrically evoked synaptic currents for l2/3 pyr neurons (left) and FSIs (right) **f,** Schematic representation of observed changes onto l2/3 pyr neurons and FSIs through analysis of excitatory and inhibitory postsynaptic currents (EPSCs & IPSCs, respectively).

## Supplementary Table

**Supplementary Table 1:** Stages of acquisition for the 2-armed bandit task (2-ABT) Center poke followed by a side poke behavior is shaped over the course of 6 stages where experimenter shifts the amount of time to make center to side selection (‘decision window’ in seconds), probability of the highly rewarded (p(high port)) and lowly rewarded (p(low port)) ports and the length of trials that the rule is held constant (‘blocklength’). Stage 7 represents the final stage of the 2-ABT that is used for the execution phase.

| stage | decision window | p(high port) | p(low port) | blocklength |
| --- | --- | --- | --- | --- |
| 1 | NA | 1.0 | 1.0 | NA |
| 2 | 15 | 1.0 | 0.0 | 50 |
| 3 | 7 | 1.0 | 0.0 | 50 |
| 4 | 3 | 1.0 | 0.0 | 50 |
| 5 | 2 | 1.0 | 0.0 | 30 |
| 6 | 2 | 1.0 | 0.0 | 20 |
| 7 | 2 | 0.8 | 0.2 | 10-30 |

## References

1. Spear, L.P., The adolescent brain and age-related behavioral manifestations. Neuroscience & biobehavioral reviews, 2000. 24(4): p. 417–463.

2. Tervo-Clemmens, B., et al., A canonical trajectory of executive function maturation from adolescence to adulthood. Nature communications, 2023. 14(1): p. 6922.

3. Larsen, B. and B. Luna, Adolescence as a neurobiological critical period for the development of higher-order cognition. Neurosci. Biobehav. Rev., 2018. 94: p. 179–195.

4. Luna, B., A. Padmanabhan, and K. O’Hearn, What has fMRI told us about the development of cognitive control through adolescence? Brain Cogn., 2010. 72(1): p. 101–113.

5. Miller, D.J., et al., Prolonged myelination in human neocortical evolution. Proc Natl Acad Sci U S A, 2012. 109(41): p. 16480–5.

6. Giorgio, A., et al., Longitudinal changes in grey and white matter during adolescence. Neuroimage, 2010. 49(1): p. 94–103.

7. Huttenlocher, P.R., Synaptic density in human frontal cortex - developmental changes and effects of aging. Brain Res, 1979. 163(2): p. 195–205.

8. Petanjek, Z., et al., The Protracted Maturation of Associative Layer IIIC Pyramidal Neurons in the Human Prefrontal Cortex During Childhood: A Major Role in Cognitive Development and Selective Alteration in Autism. Front Psychiatry, 2019. 10: p. 122.

9. Candelaria-Cook, F.T., et al., Developmental trajectory of MEG resting-state oscillatory activity in children and adolescents: a longitudinal reliability study. Cerebral Cortex, 2022. 32(23): p. 5404–5419.

10. Liuzzi, L., et al., Changes in behavior and neural dynamics across adolescent development. J. Neurosci., 2023. 43(50): p. 8723–8732.

11. Whitford, T.J., et al., Brain maturation in adolescence: concurrent changes in neuroanatomy and neurophysiology. Hum. Brain Mapp., 2007. 28(3): p. 228–237.

12. Calabro, F.J., et al., Development of hippocampal–prefrontal cortex interactions through adolescence. Cerebral Cortex, 2020. 30(3): p. 1548–1558.

13. Tang, L., A.T. Shafer, and N. Ofen, Prefrontal cortex contributions to the development of memory formation. Cerebral Cortex, 2018. 28(9): p. 3295–3308.

14. Cromer, J.A., et al., The nature and rate of cognitive maturation from late childhood to adulthood. Frontiers in psychology, 2015. 6: p. 127486.

15. Luciana, M., et al., The development of nonverbal working memory and executive control processes in adolescents. Child development, 2005. 76(3): p. 697–712.

16. Luna, B., et al., Maturation of cognitive processes from late childhood to adulthood. Child development, 2004. 75(5): p. 1357–1372.

17. Ordaz, S.J., et al., Longitudinal growth curves of brain function underlying inhibitory control through adolescence. Journal of Neuroscience, 2013. 33(46): p. 18109–18124.

18. Quach, A., et al., Adolescent development of inhibitory control and substance use vulnerability: A longitudinal neuroimaging study. Developmental cognitive neuroscience, 2020. 42: p. 100771.

19. Flynn, S.W., et al., Abnormalities of myelination in schizophrenia detected in vivo with MRI, and post-mortem with analysis of oligodendrocyte proteins. Mol Psychiatry, 2003. 8(9): p. 811–20.

20. Glantz, L.A. and D.A. Lewis, Reduction of synaptophysin immunoreactivity in the prefrontal cortex of subjects with schizophrenia. Regional and diagnostic specificity. Arch Gen Psychiatry, 1997. 54(10): p. 943–52.

21. Glantz, L.A. and D.A. Lewis, Dendritic spine density in schizophrenia and depression. Arch Gen Psychiatry, 2001. 58(2): p. 203.

22. Goto, Y., C.R. Yang, and S. Otani, Functional and dysfunctional synaptic plasticity in prefrontal cortex: roles in psychiatric disorders. Biological psychiatry, 2010. 67(3): p. 199–207.

23. Kumari, V., et al., Cortical grey matter volume and sensorimotor gating in schizophrenia. Cortex, 2008. 44(9): p. 1206–14.

24. Li, M., C. Long, and L. Yang, Hippocampal-prefrontal circuit and disrupted functional connectivity in psychiatric and neurodegenerative disorders. BioMed research international, 2015. 2015.

25. Narayanan, M., et al., Common dysregulation network in the human prefrontal cortex underlies two neurodegenerative diseases. Molecular systems biology, 2014. 10(7): p. 743.

26. Kroon, T., et al., Early postnatal development of pyramidal neurons across layers of the mouse medial prefrontal cortex. Sci Rep, 2019. 9(1): p. 5037.

27. Caballero, A., et al., Differential regulation of parvalbumin and calretinin interneurons in the prefrontal cortex during adolescence. Brain Structure and Function, 2014. 219: p. 395–406.

28. Caballero, A., et al., Emergence of GABAergic-dependent regulation of input-specific plasticity in the adult rat prefrontal cortex during adolescence. Psychopharmacology, 2014. 231: p. 1789–1796.

29. Bissonette, G.B., et al., Double dissociation of the effects of medial and orbital prefrontal cortical lesions on attentional and affective shifts in mice. Journal of Neuroscience, 2008. 28(44): p. 11124–11130.

30. Benoit, L.J., et al., Adolescent thalamic inhibition leads to long-lasting impairments in prefrontal cortex function. Nature neuroscience, 2022. 25(6): p. 714–725.

31. Ferguson, B.R. and W.-J. Gao, Thalamic control of cognition and social behavior via regulation of gamma-aminobutyric acidergic signaling and excitation/inhibition balance in the medial prefrontal cortex. Biological psychiatry, 2018. 83(8): p. 657–669.

32. Canetta, S.E., et al., Mature parvalbumin interneuron function in prefrontal cortex requires activity during a postnatal sensitive period. Elife, 2022. 11: p. e80324.

33. Laubach, M., et al., *What, if anything, is rodent prefrontal cortex?* eneuro, 2018. 5(5).

34. Le Merre, P., S. Ährlund-Richter, and M. Carlén, The mouse prefrontal cortex: Unity in diversity. Neuron, 2021. 109(12): p. 1925–1944.

35. Ferguson, B.R. and W.-J. Gao, Development of thalamocortical connections between the mediodorsal thalamus and the prefrontal cortex and its implication in cognition. Frontiers in human neuroscience, 2015. 8: p. 1027.

36. Ouhaz, Z., et al., Mediodorsal thalamus is critical for updating during extradimensional shifts but not reversals in the attentional set-shifting task. Eneuro, 2022. 9(2).

37. Le Magueresse, C. and H. Monyer, GABAergic interneurons shape the functional maturation of the cortex. Neuron, 2013. 77(3): p. 388–405.

38. Reh, R.K., et al., Critical period regulation across multiple timescales. Proc. Natl. Acad. Sci. U. S. A., 2020. 117(38): p. 23242–23251.

39. Hensch, T.K., Critical period plasticity in local cortical circuits. Nat. Rev. Neurosci., 2005. 6(11): p. 877–888.

40. Takesian, A.E. and T.K. Hensch, Balancing plasticity/stability across brain development. Prog. Brain Res., 2013. 207: p. 3–34.

41. Mukherjee, A., et al., Long-lasting rescue of network and cognitive dysfunction in a genetic schizophrenia model. Cell, 2019. 178(6): p. 1387–1402. e14.

42. Beron, C.C., et al., Mice exhibit stochastic and efficient action switching during probabilistic decision making. Proc Natl Acad Sci U S A, 2022. 119(15): p. e2113961119.

43. Vardalaki, D., K. Chung, and M.T. Harnett, Filopodia are a structural substrate for silent synapses in adult neocortex. Nature, 2022. 612(7939): p. 323–327.

44. Little, J.P. and A.G. Carter, Subcellular synaptic connectivity of layer 2 pyramidal neurons in the medial prefrontal cortex. J Neurosci, 2012. 32(37): p. 12808–19.

45. Abitz, M., et al., Excess of neurons in the human newborn mediodorsal thalamus compared with that of the adult. Cerebral Cortex, 2007. 17(11): p. 2573–2578.

46. Rios, O. and J. Villalobos, Postnatal development of the afferent projections from the dorsomedial thalamic nucleus to the frontal cortex in mice. Developmental brain research, 2004. 150(1): p. 47–50.

47. Tuncdemir, S.N., et al., Early somatostatin interneuron connectivity mediates the maturation of deep layer cortical circuits. Neuron, 2016. 89(3): p. 521–535.

48. Takesian, A.E., V.C. Kotak, and D.H. Sanes, Presynaptic GABA(B) receptors regulate experience-dependent development of inhibitory short-term plasticity. J. Neurosci., 2010. 30(7): p. 2716–2727.

49. Bartley, A.F., et al., Differential activity-dependent, homeostatic plasticity of two neocortical inhibitory circuits. J. Neurophysiol., 2008. 100(4): p. 1983–1994.

50. Miao, Q., et al., Selective maturation of temporal dynamics of intracortical excitatory transmission at the critical period onset. Cell Rep., 2016. 16(6): p. 1677–1689.

51. McKeon, S.D., et al., *Prefrontal excitation/ inhibition balance supports adolescent enhancements in circuit signal to noise ratio.* bioRxivorg, 2024.

52. Perica, M.I., et al., Development of frontal GABA and glutamate supports excitation/inhibition balance from adolescence into adulthood. Prog. Neurobiol., 2022. 219(102370): p. 102370.

53. Xue, M., B.V. Atallah, and M. Scanziani, Equalizing excitation–inhibition ratios across visual cortical neurons. Nature, 2014. 511(7511): p. 596–600.

54. Shoji, H., et al., Age-related changes in behavior in C57BL/6J mice from young adulthood to middle age. Mol. Brain, 2016. 9: p. 11.

55. Baum, M.L., et al., *CSMD1 regulates brain complement activity and circuit development.* Brain, Behavior, and Immunity, 2024. 119: p. 317–332.

56. Sekar, A., et al., Schizophrenia risk from complex variation of complement component 4. Nature, 2016. 530(7589): p. 177–183.

57. Singh, T., et al., Rare coding variants in ten genes confer substantial risk for schizophrenia. Nature, 2022. 604(7906): p. 509–516.

58. Vormstein-Schneider, D., et al., Viral manipulation of functionally distinct interneurons in mice, non-human primates and humans. Nat Neurosci, 2020. 23(12): p. 1629–1636.

59. Chantranupong, L., et al., Dopamine and glutamate regulate striatal acetylcholine in decision-making. Nature, 2023. 621(7979): p. 577–585.

60. Merel, J., et al., Bayesian methods for event analysis of intracellular currents. J Neurosci Methods, 2016. 269: p. 21–32.

61. Wilton, D.K., et al., Microglia and complement mediate early corticostriatal synapse loss and cognitive dysfunction in Huntington’s disease. Nat Med, 2023. 29(11): p. 2866–2884.

62. Antoine, M.W., et al., Increased Excitation-Inhibition Ratio Stabilizes Synapse and Circuit Excitability in Four Autism Mouse Models. Neuron, 2019. 101(4): p. 648–661 e4.

63. Nagai, Y., et al., Deschloroclozapine, a potent and selective chemogenetic actuator enables rapid neuronal and behavioral modulations in mice and monkeys. Nat Neurosci, 2020. 23(9): p. 1157–1167.

